# Paradoxically sparse chemosensory tuning in broadly-integrating external granule cells in the mouse accessory olfactory bulb

**DOI:** 10.1101/703892

**Authors:** Xingjian Zhang, Julian P. Meeks

## Abstract

The accessory olfactory bulb (AOB) is a critical circuit in the mouse accessory olfactory system (AOS), but AOB processing is poorly understood compared to the main olfactory bulb (MOB). We used 2-photon GCaMP6f Ca^2+^ imaging in an *ex vivo* preparation to study the chemosensory tuning of AOB external granule cells (EGCs), an interneuron population hypothesized to broadly integrate from mitral cells (MCs). We measured MC and EGC tuning to natural chemosignal blends and monomolecular ligands, finding that EGC tuning was far sparser than MC tuning. Simultaneous patch-clamp electrophysiology and Ca^2+^ imaging indicated that this was only partially explained by lower GCaMP6f-to-spiking ratios in EGCs compared to MCs. *Ex vivo* patch-clamp recordings revealed that EGC subthreshold responsivity was broad, but monomolecular ligand responses were insufficient to elicit spiking. These results indicate that EGC spiking is selectively engaged by chemosensory blends, suggesting different roles for EGCs than analogous interneurons in the MOB.

## Introduction

Social behavior involves multimodal sensory inputs, information processing cascades carried out by multilevel neural circuits, and complicated behavioral outputs. Within each brain region, interactions between principal neurons and interneurons form neural circuit motifs, the fundamental computational building blocks for information processing (Braganza and Beck, 2018). In mice and many mammals, the accessory olfactory system (AOS) is required for the expression of typical social behaviors, but many fundamentals of its neural circuit organization and function remain unclear. The AOS detects non-volatile chemosignals including pheromones (for intra-species communication) and kairomones (for inter-species communication) (Su et al., 2009). In rodents, the first dedicated neural circuit for chemosignal information processing is the accessory olfactory bulb (AOB).

AOB principal neurons, known as mitral cells (MC), are tightly modulated by local GABAergic interneurons and GABAergic modulation in the AOB has significant behavioral impacts. (Kaba and Nakanishi, 1995; Brennan, 2001; Hendrickson et al., 2008). For example, microinfusion of GABAergic transmission blocker to AOB can block pregnancy (Kaba and Keverne, 1988). Many studies have implicated AOB GABAergic interneurons in social behaviors and experience-dependent chemosensory plasticity (Brennan et al., 1995; Matsuoka et al., 1997; Matsuoka et al., 2004; Brennan and Binns, 2005; Kaba and Huang, 2005; Araneda and Firestein, 2006; Hendrickson et al., 2008; Cansler et al., 2017; Gao et al., 2017). Cumulatively, these observations suggest that MC-interneuron communication in the AOB is critical for rodent physiology and behavior.

Despite recent progress, many gaps in our understanding of AOB MC-interneuron interactions remain. A major limitation for our understanding of AOB function is a lack of information about how MCs interact with several inhibitory neural types. The AOB has a variety of interneurons that have been largely classified based on their location and morphology (Larriva-Sahd, 2008; Moriya-Ito et al., 2013). Three major AOB interneuron types have been identified, along with several minor types. Juxtaglomerular cells (JGCs) are found within the superficial glomerular layer, where excitatory sensory inputs from the vomeronasal organ (VNO) enter the AOB. External granule cells (EGC) are found in the external cell layer alongside the somas of MCs. Internal granule cells (IGCs), the most abundant and well-studied AOB interneuron type, are located in the internal granule layer. JGCs, similar to counterparts in the main olfactory bulb (MOB), are thought to modulate the input to the MCs via their synaptic connection with VNO axon terminals and MC apical dendrites within glomeruli (Geramita and Urban, 2017). EGCs and IGCs, in contrast, are thought to modulate MC activity via reciprocal dendrodendritic connections (Taniguchi and Kaba, 2001; Castro et al., 2007). Relative to JGCs and IGCs, there is very little information about the physiology or function of AOB EGCs; just one targeted study of their intrinsic features has been reported (Maksimova et al., 2019). Cells that appear analogous to AOB EGCs have been studied in the MOB. Specifically, MOB parvalbumin-expressing interneurons in the external plexiform layer (PV-EPL interneurons) resemble EGCs in their morphologies and apparent broad connectivity with MCs. These MOB PV-EPL interneurons were shown to integrate excitatory information from multiple MCs, resulting in broader odorant receptive fields than MCs (Kato et al., 2013). The broad tuning of MOB PV-EPL interneurons underscored a role these cells in divisive normalization of MCs via lateral feedback inhibition (Kato et al., 2013; Miyamichi et al., 2013). Given the similarities between MOB PV-EPL interneurons and AOB EGCs, we hypothesized that AOB EGCs would display broad chemosensory tuning and carry out divisive normalization of MCs through their inhibitory dendro-dendritic synapses.

Here, we describe the first targeted investigation of AOB EGCs in the context of chemosensory function. We studied EGC function using two-photon GCaMP6f Ca^2+^ imaging and patch clamp electrophysiology in a specialized *ex vivo* preparation that preserves functional connectivity between the VNO and AOB (Meeks and Holy, 2009). This preparation enabled direct observation of sensory integration performed by AOB MCs during peripheral stimulation with known AOS activators, including monomolecular steroid ligands and natural ligand blends (mouse urine and feces). Using a Cre-expressing transgenic mouse (*Cort*-cre), which selectively labels subsets of AOB EGCs (Taniguchi et al., 2011; Maksimova et al., 2019), we measured EGC activation by AOS ligands, finding unexpectedly sparse activation compared to MCs and JGCs. Because this observation was at odds with our expectations, we performed whole-cell patch clamp experiments on EGCs, finding that GCaMP6f signals in EGCs weakly report spiking activity. Furthermore, we found that EGCs indeed broadly integrate from MCs, but EGCs rarely fire action potentials unless a natural blend containing many chemosensory cues (*e.g.* mouse urine or feces) was used to stimulate the VNO. These results indicate that AOB EGCs modulate the circuit activity in a fashion unlike their MOB counterparts. The lack of broadly tuned spiking activity does not support divisive normalization in the context of small numbers of odorants. Instead, these data suggest that EGCs inhibit MCs in specific chemosensory conditions involving rich pheromone environments or perhaps that they modulate MC activity through spiking-independent mechanisms (Isaacson and Strowbridge, 1998; Schoppa et al., 1998; Chen et al., 2000; Halabisky et al., 2000; Isaacson, 2001; Egger et al., 2005; Bywalez et al., 2015; Lage-Rupprecht et al., 2018). These studies provide new information about the role of AOB EGCs in AOS sensory processing, and place important constraints on our models of AOB circuit function.

## Materials and Methods

### Mice

All animal procedures were in compliance with the UT Southwestern Institutional Care and Use Committee. Mice used in this research were C57BL/6J unless otherwise noted. *Cort*-T2A-Cre and *Gad*-IRES-Cre (Taniguchi et al., 2011) were from The Jackson Laboratory (Stock# 010910 and 028867). *Pcdh21*-Cre (Nagai et al., 2005) mice were kindly shared by the laboratory of Timothy Holy with permission from the originating institution. Both male and female mice were used in all experiments and the results pooled. 27 mice (14 females and 13 males) were used for EGC Ca^2+^ imaging. 5 mice (3 females and 2 males) were used for JGC Ca^2+^ imaging. 9 mice (4 females and 5 males) were used for MC Ca^2+^ imaging. For EGC *ex vivo* patch clamp recording, 16 mice (10 females and 6 males) were used.

### Stimuli and reagents

Female mouse fecal extracts and urine were prepared as previously described (Nodari et al., 2008; Meeks et al., 2010; Doyle et al., 2016). Fecal extracts and urine were pooled across subjects of the same sex, strain, and age, then aliquoted and stored at −80 °C. Just prior to each experiment, aliquots were thawed and diluted in control Ringer’s saline solution containing (in mM): 115 NaCl, 5 KCl, 2 CaCl_2_, 2 MgCl_2_, 25 NaHCO_3_, 10 HEPES and 10 glucose. For VNO stimulation, the fecal extracts were diluted at 1:300 and the urine was diluted at 1:100, concentrations that activate roughly equal number of AOB MCs in the *ex vivo* preparation (Doyle et al., 2014).

All sulfated steroids were purchased from Steraloids, Inc. (Newport, RI, USA). The sulfated steroid panel includes A7864 (5-androsten-3β, 17β-diol disulphate, disodium salt), A6940 (4-androsten-17α-ol-3-one sulphate sodium salt), A7010 (4-androsten-17β-ol-3-one sulphate, sodium salt), E0893 (1, 3, 5(10)-estratrien-3, 17α-diol 3-sulphate, sodium salt), E1050 (1, 3, 5(10)-estratrien-3, 17β-diol disulphate, disodium salt), E4105 (4-estren-17β-ol-3-one sulphate, sodium salt), P3817 (5α-pregnan-3α-ol-20-one sulphate sodium salt), P3865 (5α-pregnan-3β-ol-20-one sulphate, sodium salt), P8168 (5β-pregnan-3α-ol-20-one sulphate, sodium salt), Q1570 (4-pregnen-11β, 21-diol-3, 20-dione 21-sulphate, sodium salt) and Q3910 (4-pregnen-11β, 17, 21-triol-3, 20-dione 21-sulphate, sodium salt). 20 mM stock solutions of A7864, E1050 and Q1570 were prepared in H_2_O, the 20 mM stock solution of all other sulfated steroids were prepared in methanol. Upon use, stock solutions were diluted at 1:2000 into the Ringer’s solution (10 μM working concentration). Methanol was diluted at 1:2000 into the Ringer’s solution as a vehicle control.

### Virus injection

GCaMP6f expression in AOB neurons was achieved by injecting AAV.CAG.Flex.GCaMP6f.WPRE.SV40 to the corresponding Cre mouse lines. To achieve optimal GCaMP6f expression, different AAV pseudotypes were used. AAV9 (Penn Vector Core, Catalog #AV9-PV2816) was used on *Cort*-IRES-Cre for EGC labeling, and *Gad2*-IRES-Cre for JGC labeling. AAV5 (Penn Vector Core, Catalog #AV5-PV2816) was used on *Pcdh21*-Cre for MC labeling, confirming the efficacy of this AAV pseudotype for MCs (Rothermel et al., 2013).

Adult mice of 8-12 weeks old were used for virus injection. Intracranial injections were performed on a customized stereotaxic device that rotated the mouse head such that the rostral end of the head tilted up ∼30°. Mice were anesthetized via isofluorane inhalation using a SomnoSuite Small Animal Anesthesia System (Kent Scientific). For each animal, ∼180-300 nL viral vector (≥ 1e^13^ vg/ml) was injected into AOB. The bilateral coordinates, measured from the lambda, were x ∼ ±1000 μm, y ∼ 4150 μm, z ∼ 3300 μm for 8-week old mice. After virus injection, the animals were allowed to recover for at least 3 weeks before being used for experiments.

### VNO-AOB *ex vivo* preparation

*Ex vivo* preparations were performed as described previously (Meeks and Holy, 2009; Doyle et al., 2014). Briefly, mice were anesthetized by isoflurane inhalation, followed by rapid decapitation into the ice-cold aCSF. After removing the scalp, the snout and olfactory bulbs were separated from the rest of the skull, and the snout was then halved along the midline, maintaining the VNO AOB from the right hemisphere. The resulting tissue was affixed to a plastic plank with tissue adhesive (Krazy Glue, Elmer’s Products) and placed into a custom perfusion chamber where secondary dissections were performed. In this chamber, room temperature (22-25 °C) oxygenated artificial cerebrospinal fluid (aCSF) was rapidly superfused over the tissue at a rate of 5-8 mL/min. aCSF contained (in mM): 125 NaCl, 2.5 KCl, 2 CaCl_2_, 1 MgCl_2_, 25 NaHCO_3_, 1.25 NaH_2_PO_4_, 25 glucose, 3 *myo*-inositol, 2 sodium pyruvate, and 0.4 sodium ascorbate. The septal cartilage was carefully removed, exposing the septal tissue containing the axons from VNO to the oxygenated aCSF. The sample was then transferred to a second, custom-built tissue chamber with a rotatable platform. A small cut was made at the anterior end of the VNO capsule, through which polyimide tubing (A-M Systems, 0.0045’’ID, 0.00050’’ WALL) was inserted for stimulus delivery. Stimulation solution was pressurized at 9-12 psi giving an effective flow rate of 0.2-1 mL/min. Valve opening was controlled by Automate Scientific perfusion system with ValveLink8.2 controller. Once cannulated, the platform was rotated so that the AOB facing upward to facilitate 2-photon imaging on an upright microscope.

### 2-photon ex vivo GCaMP6f imaging

Adult mice aged 11-16 weeks were used for imaging. *Ex vivo* preparations in the customized chamber were placed into a custom adapter on a Thorlabs Acerra upright 2-photon microscope system equipped with an OLYMPUS XLUMPlanFLN 20X objective and a fast-scanning resonant galvanometer along one of the two principal axes. To excite GCaMP6f fluorescence, 910 nm light (average power 2100 mW measured at the laser, 25-35% power transmission for imaging) was used. Images with pixel dimensions 512×512 were acquired at 30 frames per second and synchronized with stimulus delivery system (Automate Scientific) via Axon Clampex 10 software (Molecular Devices). Episodic stimulation sessions, consisting of 1 s pre-stimulation VNO Ringer’s solution flush, 8 s of VNO stimuli, and 11 s post-stimulation VNO Ringer’s solution flush, were used to present multiple repeats per cell. Across sessions, stimulus presentation order was randomized to reduce the impact of potential stimulus order effects.

### Acute slice preparation

Mice were anesthetized with isofluorane and immediately decapitated into ice-cold oxygenated aCSF with an additional 9 mM MgCl_2_. Brains were then extracted and a vertical cut at the prefrontal cortex was made and the anterior part containing the olfactory bulbs was preserved. Another vertical cut along the midline separated the two hemispheres, and both were embedded in aCSF containing 3% low-melt agarose at 37 °C. The agarose block was then mounted on an angled slicing platform on a vibrating microtome (Leica VT1200). The slicing blade ran at an angle of approximately 12 degrees off-sagittal, running from caudal/medial to rostral/lateral. The slices were then collected in recovery chamber containing oxygenated room-temperature aCSF with 0.5 mM kynurenic acid. Slices were allowed to recover at least 30 min before being used for patch clamp recording.

### Electrophysiology

Acute slice electrophysiology was performed on the same upright 2-photon microscope used in *ex vivo* imaging. Slices were placed in a tissue chamber (Warner instruments), warmed to 28-30 °C by a temperature controller (Warner instruments). GCaMP6f-expressing neurons were identified using the same laser setup with *ex vivo* imaging. Thin borosilicate glass electrodes (TW150, World Precision Instruments) were pulled using a horizontal puller (P1000, Sutter Instruments). The electrodes were then filled with standard internal solution containing 115 mM K-gluconate, 20 mM KCl, 10 mM HEPES, 2 mM EGTA, 2 mM MgATP, 0.3 mM Na_2_GTP, 10 mM Na phosphocreatine at pH 7.37. AlexaFluor568 (166 μM, Thermo Fisher) was added for visualization under 2-photon microscope. Pipette resistance ranged from 7-10 MΩ for EGC patch clamp, and 6-8 MΩ for MC patch clamp. The electrodes were controlled with a MicroStar motorized micro-manipulator (Scientifica). In the experiments probing the relationship between GCaMP6f signal and action potentials, after whole-cell configuration was formed, cells were voltage clamped at −70 mV, and a 10 s train of 20 Hz depolarization pulses to 0 mV (2 ms pulse width) was delivered through the electrode. GCaMP6f signals were recorded simultaneously using the same imaging parameters as *ex vivo* Ca^2+^ imaging experiments. The GCaMP6f signal was then extracted using custom MATLAB programs.

The *ex vivo* preparation setup for whole-cell patch clamp experiments was the same as for the *ex vivo* imaging. *Cort*-cre mice were crossed with Cre-dependent tdTomato effector mice (“Ai9” mouse line; Jackson Laboratory stock #007909) to label *Cort*+ cells. AlexaFluor488 (100 μM, Thermo Fisher) was added in the standard internal solution for electrode visualization under 2-photon microscope. Target cells were identified and approached using the ‘approach’ mode of the micro-manipulator (to facilitate penetrating the tissue without tearing the glomerular layer) prior to achieving the whole-cell configuration. After the whole-cell configuration was achieved, cells were held in current clamp mode. Upon break-in, the resting membrane potential of the cell was measured, and steady-state holding current was applied throughout the experiment to maintain the initial resting membrane potential. The same panel of monomolecular ligands and natural stimuli were applied to the VNO as the *ex vivo* Ca^2+^ imaging experiments.

All recordings were amplified via a MultiClamp 700B amplifier (Molecular Devices) at 20 kHz and were digitized by a DigiData 1440 analog-digital converter via pClamp 10.5 software (Molecular Devices, RRID: SCR_011323). Data were analyzed by custom software written in MATLAB and graphs were created using MATLAB and R (ggplot2).

## Data analysis

### 2-photon ex vivo GCaMP6f imaging

Raw 2-photon Ca^2+^ imaging analysis was performed using customized MATLAB scripts. ROIs were manually selected and ΔF/F values were extracted by comparing the change in fluorescence during stimulation to 30 frames (∼1 s) prior to each stimulus session. Further analysis of ΔF/F signals was performed using customized R scripts and graphs were made using ggplot2. ΔF/F responses were averaged over 4 or more trials for each stimulus. Because the latency to peak for each cell and each specific preparation can vary (typically between 7s and 12 s from stimulation onset) we used average response curves to determine a continuous series of samples during which the ΔF/F value is above 50% of the peak value. For each individual repeat, we integrated the ΔF/F intensity during this time window. We assessed the statistical reliability of each cell’s stimulus responsiveness using the unpaired Student’s *t*-test, comparing each stimulus to the vehicle control trials. We considered a cell to be responsive to a stimulus if (1) its p-value was less than 0.05 and peak ΔF/F was greater than 0.1 or (2) its p-value was less than 0.1 and peak ΔF/F greater than 0.3). For heat map displays throughout the manuscript, average peak ΔF/F value was used to represent each cell’s response strength towards to each stimulation. Cumulative distribution of EGCs, JGCs and MCs tuning were evaluated using Kolmogorov–Smirnov test.

### 2-photon ex vivo whole cell patch clamp

In these experiments, each round of stimulation included 1 s pre-stimulation flush, 8 s stimulation and 11 s post-stimulation flush. The stimulation panel was split into 2 bouts, each consisting of 6 sulfated steroids, 2 naturalistic stimuli, and the vehicle control, delivered in randomized orders. 2 bouts were used to cover the entire stimulation panel, and at least X full repeats of the full stimulus panel were used for all analyzed experiments. The membrane voltage was recorded at 20 kHz during the stimulation administration, and for all comparisons except spike analysis the data was downsampled by decimation by a factor of 100. For each repeat we calculated the average value of the top 5% of voltage reads in a static time window between 1.5 and 10 s following the stimulus onset and used the value to quantify the subthreshold activity. The response to each stimulus was compared to the vehicle control using the Wilcoxon rank sum test (4 or more repeats per stimulus). All stimulus responses with p < 0.05 were considered effective.

## Results

### Implementation of cell type-specific GCaMP6f Ca^2+^ imaging in the AOB ex vivo

The mouse AOB remains one of the most poorly understood principal sensory circuits in the mammalian brain. A large reason for this deficiency is the limited number of studies on the sensory responses of AOB neurons. Several *in vivo* and *ex vivo* studies have investigated MC sensory responses, but studies of interneuron function are severely lacking (Luo et al., 2003; Hendrickson et al., 2008; Ben-Shaul et al., 2010; Meeks et al., 2010; Doyle et al., 2016). We used combined *ex vivo* sensory preparations that retain VNO-AOB connectivity (Meeks and Holy, 2009) with 2-photon GCaMP6f imaging to measure Ca^2+^ signals in specific AOB neuronal populations (Fig. 1A).

**Figure 1.**
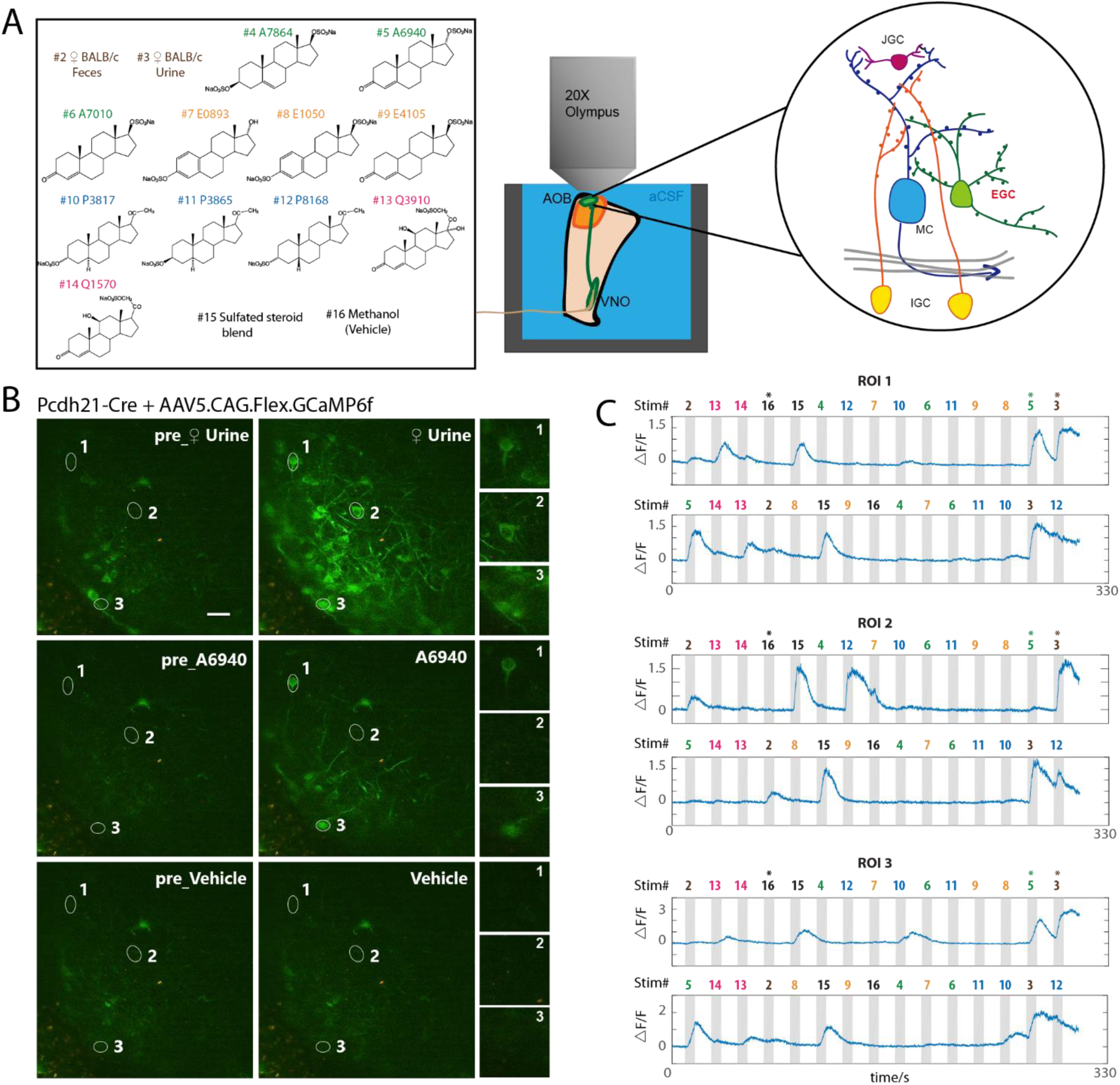
(**A**). Overview of e*x vivo* Ca^2+^ imaging. Left: the stimulus panel delivered to the VNO to drive activity in the AOB. Included are natural ligand blends (1:100 diluted BALB/c mouse urine and 1:300 diluted mouse feces) and 11 monomolecular sulfated steroids at 10 µM. Right: diagram of AOB circuit. MC: mitral cell, EGC: external granule cell, JGC: juxtaglomerular cell, IGC: internal granule cell. (**B**). Raw images of GCaMP6f fluorescence during *ex vivo* Ca^2+^ imaging experiments on AOB MCs. MCs expressed GCaMP6f via the infusion of Cre-dependent AAVs into the AOBs of *Pcdh21*-Cre transgenic mice 3 or more weeks prior to the recordings. Numbered regions of interest denote 3 highlighted MCs with different tuning preferences. (**C**). ΔF/F measurements from the 3 cells highlighted in (B) across 2 randomized repeats. Numbers above the gray vertical bars indicate the stimulus being applied, with colors matched to the stimulus panel in (A).

An important consideration for any study of chemosensory tuning is that measured receptive fields critically depend on the choice of chemosensory cues and concentrations. Some physiological studies of AOB tuning have exclusively utilized natural blends of chemosensory cues (e.g. dilute urine and saliva)(Hendrickson et al., 2008; Ben-Shaul et al., 2010; Tolokh et al., 2013), whereas others have used both of natural chemosignal blends and monomolecular VNO ligands (Meeks et al., 2010; Doyle et al., 2016; Doyle and Meeks, 2017). We chose to use both natural and monomolecular stimuli; we selected a panel that included diluted mouse urine and fecal extracts and monomolecular sulfated steroid ligands similar to those used by previous studies (Meeks et al., 2010; Turaga and Holy, 2012; Doyle et al., 2016). We first recorded sensory tuning to this panel of odorants in AOB MCs, by virally or transgenically driving GCaMP6f in *Pcdh21*-cre transgenic mice (Nagai et al., 2005), we observed reliable, time-locked, stimulus-driven chemosensory activity in populations of AOB MCs across multiple stimulus trials (Fig. 1B, C, Supplementary Video 1). GCaMP6f responses to 8 s stimulus trials were large in amplitude (∼0.4 - ∼3.2 peak amplitude) and slow to peak and decay (peak time 7 to 12 s from stimulation onset; decay time 8 to 14 s), consistent with the time course of action potential firing observed in MCs with similar stimulation conditions (Hendrickson et al., 2008; Meeks and Holy, 2009). The establishment of GCaMP6f 2-photon Ca^2+^ imaging in the *ex vivo* preparation allowed us to investigate sensory tuning properties of genetically-defined AOB cell types.

### Mitral cell GCaMP6f imaging confirms broad chemosensory integration

AOB interneurons are principally excited by glutamatergic sensory input from AOB MCs (Brennan and Keverne, 1997; Taniguchi and Kaba, 2001). The tuning of AOB MCs to a similar panel of chemosensory stimuli has been characterized using extracellular single-unit recordings (Meeks et al., 2010). However, since the GCaMP6f imaging platform represents a new approach, we first wanted to investigate the tuning properties of genetically-defined MCs and compare these measurements to previous results (Fig. 2). Virally-driven GCaMP6f fluorescence in *Pcdh21*-cre mice was observed in cell bodies and apical dendrites of MCs spreading through the ECL and the glomerular layer (see Fig. 1B). We focused our recordings on GCaMP6f positive somas, which were all located below the AOB glomerular layer (>70 µm from the AOB surface).

**Figure 2.**
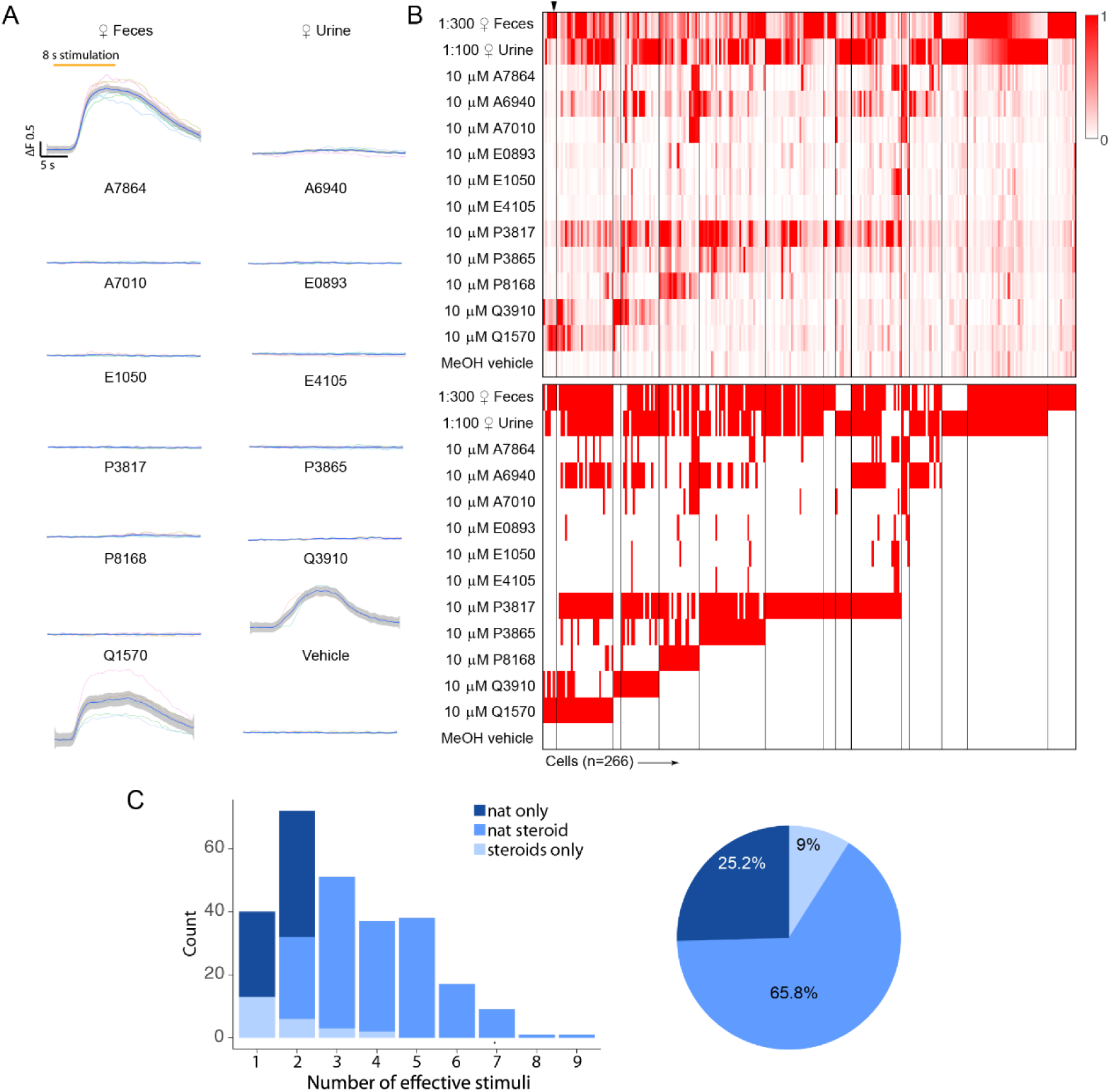
Chemosensory tuning of MCs. (**A**). Averaged response traces from an example MC. Traces were smoothed by local polynomial regression fitting. The shaded regions represent 95% confidence intervals. (**B**). Upper panel: heat map plot of normalized ΔF/F for 266 MCs. Lower panel: binary heat map plot of MC responsiveness; red tiles indicate a stimulus response that passed statistical criteria. (**C**). Left: histogram showing the number of effective stimuli per cell. Recorded cells are classified into 3 color coded groups based on their responsivity. Right: pie chart showing the composition of recorded MC populations.

We recorded chemosensory activities of 266 AOB MCs (Fig. 2B), a cohort more than 2-fold larger than previous electrophysiological studies (Hendrickson et al., 2008; Ben-Shaul et al., 2010; Meeks et al., 2010; Tolokh et al., 2013). Consistent with previous results, dilute female mouse urine and feces stimulated strong global activity that began soon after stimulus delivery. Stimulation-evoked ΔF/F typically reached a peak within the first 2 seconds of an 8 s VNO stimulus delivery, and displayed slow decay kinetics (decay time 8 – 12 s after the peak). Because the decay kinetics of GCaMP6f (Chen et al., 2013) are much faster than previous measurements of spike frequency decay, the slowness of GCaMP6f offset times likely reflects the slow cessation of spiking activity in AOB MCs (Luo et al., 2003; Wagner et al., 2006; Hendrickson et al., 2008; Ben-Shaul et al., 2010; Meeks et al., 2010; Mohrhardt et al., 2018). Of the 266 MCs we studied, 242 (91.0%) responded to at least 1 of the naturalistic stimuli, 199 (74.8%) responded to at least 1 monomolecular sulfated steroid ligand, and 125 (47.0%) were responsive to at least 2 sulfated steroids (Fig. 2A). We also observed a substantial number (24 of 266, 9%) of Pcdh21+ cells that were exclusively responsive to one or more sulfated steroids (Fig. 2C). Cluster analysis of MC stimulus responses revealed stereotyped patterns of steroid sensitivity that were consistent with previous spiking-based measurements, suggesting that MC GCaMP6f measurements accurately reflect MC activity (Fig. 2C)(Meeks et al., 2010).

### AOB juxtaglomerular cells show a slight bias towards naturalistic stimuli

AOB MCs activity is shaped at multiple levels by inhibitory interneurons. The first stage of MC inhibition occurs in the glomerular layer, where AOB JGCs reside and release GABA onto MC dendrites and VSN presynaptic terminals (Mohrhardt et al., 2018). Because there have not been any systematic recordings of AOB interneuron tuning, we first sought to measure tuning in a general population of AOB GABAergic interneurons. We therefore expressed GCaMP6f in AOB interneurons by stereotaxically injecting AAV9.CAG.Flex.GCaMP6f into the AOB of *Gad2*-IRES-Cre transgenic mice (Taniguchi et al., 2011). A large population of neurons and dendritic arbors in the AOB glomerular layer and the superficial external cellular layer were strongly labeled and visible under the 2-photon microscope (Supplementary Video 2). The density of GCaMP6f labeling in the deeper ECL, where EGC somas and IGC dendrites reside, paradoxically precluded the identification of well-resolved neuronal recordings. However, neurons in the GL and superficial ECL were readily observed that had small soma size (∼10 µm) and compact dendrites that ramified within the glomerular layer, consistent with anatomical descriptions of JGCs (Larriva-Sahd, 2008).

JGCs reside in the glomerular layer and sense glutamate released by VSN axons and MC dendrites (Jia et al., 1999; Castro et al., 2007). As with MCs, following VNO chemosensory stimulation we observed large, reliable GCaMP6f responses over multiple randomized trials (Supplementary Video 2). Dilute BALB/c feces and urine were the two most potent JGC activators. 198 of 203 (97.5%) recorded JGCs showed reliable response towards female mouse feces or urine (Fig. 3B, C). In the JGC dataset, 300-fold diluted female mouse feces triggered stronger global activity than 100-fold diluted female mouse urine (Fig. 3B), which may at least be a partial consequence of the most accessible imaging region being in the lateral/anterior quadrant of the AOB. Of 113 (55.7%) sulfated steroid-responsive JGCs, 93 (45.8%) were responsive to no more than 2 different sulfated steroids and 9 (4.4%) were responsive to more than 4 sulfated steroids (Fig. 3C). The response patterns of AOB JGCs to this panel were largely similar to MCs, but a slightly lower percentage of JGCs responded to both naturalistic stimuli and monomolecular ligands than AOB MCs (Fig. 2C, Fig. 3C), suggesting that, as expected based on their restricted glomerular innervation patterns, JGCs perform less excitatory integration than MCs.

**Figure 3.**
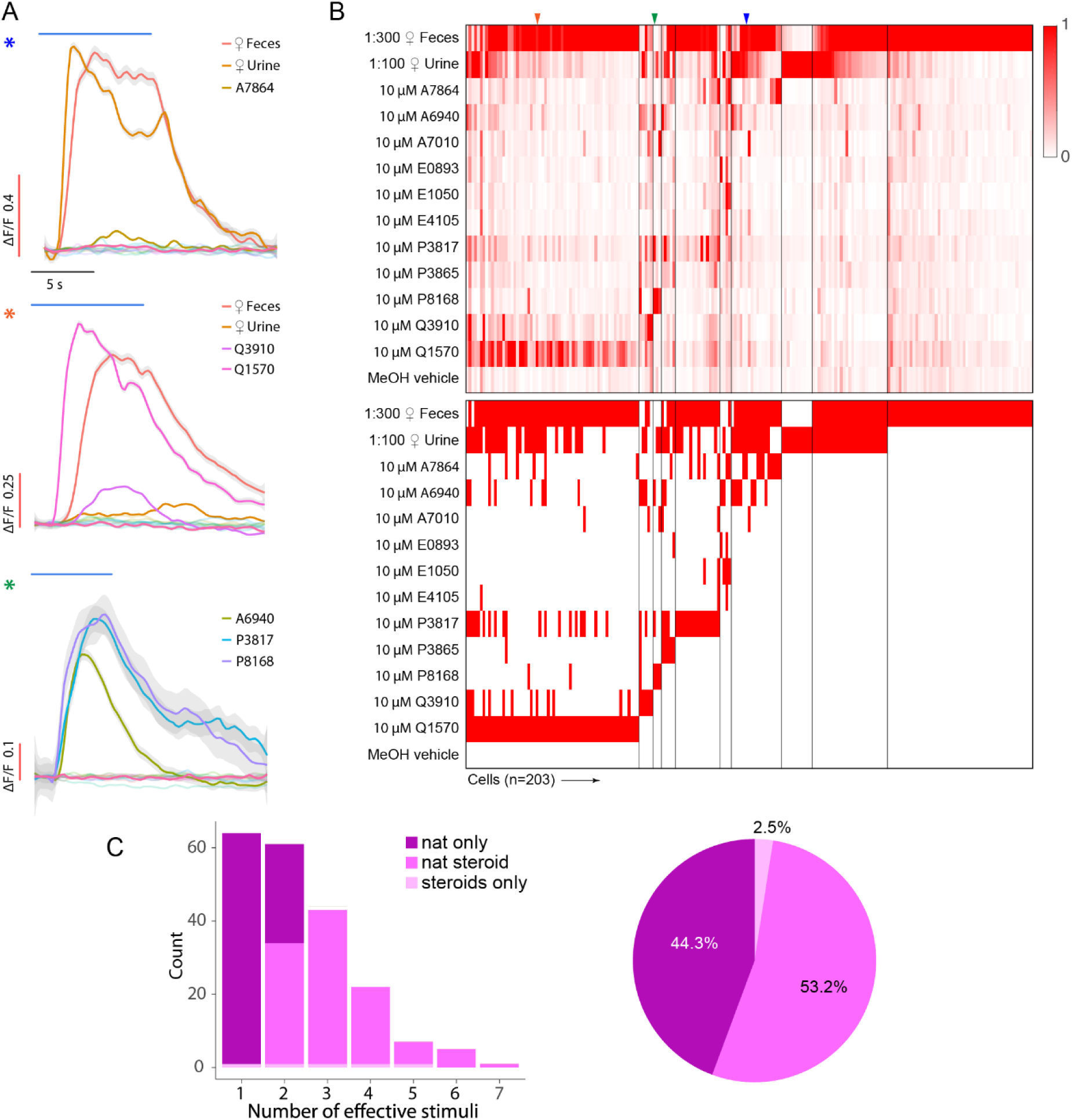
(**A**). Averaged ΔF/F traces for 3 example JGCs. Traces were smoothed by local polynomial regression fitting. The shaded regions indicate 95% confidence intervals. (**B**). Upper panel: heat map of normalized ΔF/F for 203 JGCs. Lower panel: heat map of JGC stimulus responsiveness. (**C**). Left: histogram showing the number of effective stimuli per cell. Recorded cells were classified into 3 color coded groups by their selectiveness. Right: pie chart showing the composition of recorded JGC responsivities.

### EGC GCaMP6f imaging indicates remarkably sparse chemosensory tuning

In primary chemosensory circuits, a common inhibitory motif involves broadly-integrating interneurons that perform divisive normalization or gain scaling (Wilson et al., 2012; Kato et al., 2013; Jeanne and Wilson, 2015). In the MOB, PV-EPL interneurons have been shown to perform these functions, but it is unknown whether analogous cells exist in the AOB. EGCs seemed well-matched to the morphological and physiological features of MOB PV-EPL neurons (Larriva-Sahd, 2008; Maksimova et al., 2019). Many EGCs are labeled in *Cort*-cre transgenic mice (Maksimova et al., 2019), a fact which we exploited here to specifically target EGCs for viral infection (Fig. 4A). We drove GCaMP6f expression in *Cort+* EGCs via AAV9.CAG.Flex.GCaMP6f injection into the AOB, followed by 2-photon Ca^2+^ imaging in *ex vivo* preparations.

**Figure 4.**
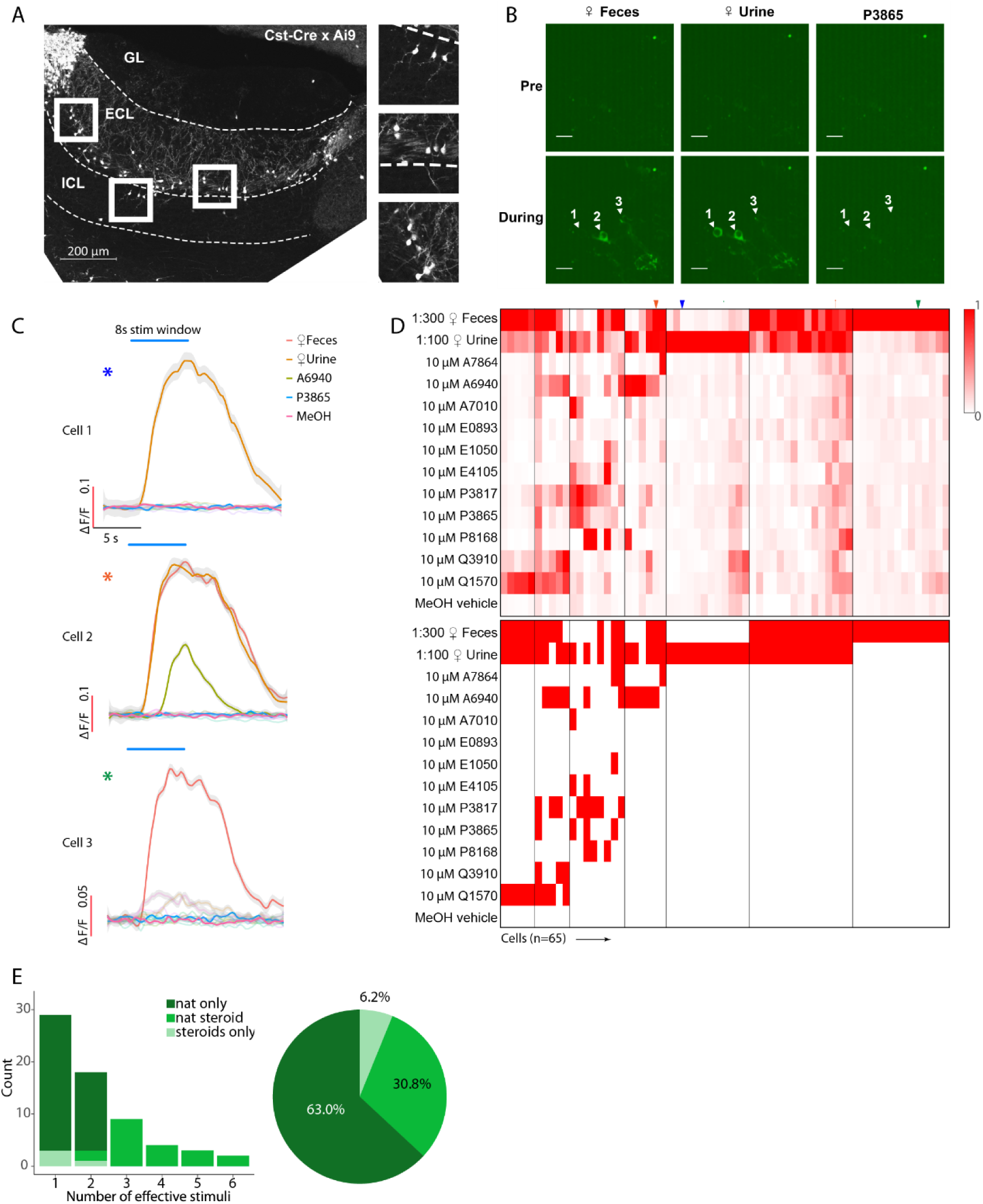
(**A**) AOB Sagittal section from a *Cort*-cre transgenic mouse mated to a cre-dependent tdTomato reporter line. (**B**). Raw GCaMP6f images from 3 example EGCs. (**C**). Averaged ΔF/F traces of the 3 cells shown in B. Traces were smoothed by local polynomial regression fitting. Shaded regions represent 95% confidence intervals. (**D**). Upper panel: heat map of normalized ΔF/F for 65 recorded EGCs. Lower panel: binary heat map of EGC stimulus responsiveness; red tiles represent the pairs that passed the statistical criteria. (**E**). Left: Histogram of EGC responsivity, with tuning classified into 3 color-coded subgroups. Right: Pie chart of EGC response classes.

GCaMP6f expression was concentrated in the AOB ECL (> 80 um from the AOB surface), but basal fluorescence intensity was extremely low compared to AOB cells infected with AAVs in both *Pcdh21*-cre and *Gad2*-cre mice (principal neurons and JGCs, respectively). Baseline GCaMP6f intensity was so low in most infected EGCs that they were not detectable above background until the AOB was activated by VNO stimulation (Fig. 4B). Identified GCaMP6f-expressing cells had small soma size, unipolar or bipolar arborizations, and in cases where GCaMP6f fluorescence was extremely high (presumably due to loss of membrane integrity/cell death), we observed large arborizations dense in synaptic spines or gemmules, all of which were consistent with previous descriptions of EGCs (Larriva-Sahd, 2008; Maksimova et al., 2019).

We next investigated the chemosensory tuning properties of Cort+ EGCs. To our surprise, these EGCs showed evidence of extremely sparse, rather than broad, tuning to the panel of chemosensory cues (Fig. 4D, E, Supplementary Video 3). Of the 65 recorded EGCs, 41 (63.0%) were activated only by naturalistic stimulation but not by sulfated steroids. Just 24 (37.0%) out of 65 Cort+ EGCs were responsive to sulfated steroids at all, with only 4 (6.2%) of the recorded cells exclusively activated by sulfated steroids (Fig 4D, E). This small population of cells showed broad sulfated steroid tuning, but also showed above-normal baseline fluorescence and spontaneous activity, perhaps suggesting that these cells may have been unhealthy (or are perhaps members of a rare Cort+ cell subtype). These experiments indicated that Cort*+* EGCs have unique features that keep basal GCaMP6f fluorescence low and revealed that EGCs are sparsely tuned to chemical cues, which was contrary to our hypothesis that they, like PV-EPL interneurons in the MOB, are more broadly tuned than their upstream MC inputs.

### Chemosensory tuning comparisons between MCs, JGCs and EGCs

To more directly investigate cell type-specific tuning in the AOB, we assessed tuning breadth to all stimuli and monomolecular steroids across all of the cell types studied. (see Methods; Fig. 5). The distributions of effective stimuli with (Fig. 5A) or without (Fig. 5B) naturalistic stimuli per cell indicated that MCs demonstrated the broadest tuning to this panel of chemosensory cues, with 103 of 266 (38.7%) being responsive to no less than 4 stimuli (Fig. 5A), and 125 of 266 (47.0%) are responsive to no less than 3 sulfated steroids. In contrast, the majority of EGCs (47 of 65, 72.3%) were responsive to 2 or fewer stimuli and 41 of 65 (43.1%) were not responsive to any sulfated steroids tested. Gad2+ JGCs demonstrated intermediate tuning in both with and without naturalistic stimuli cases. The broadness of MC tuning is consistent with heterotypic integration of VNO inputs by AOB MCs (Wagner et al., 2006; Meeks et al., 2010). When natural ligand blends were included, the cumulative distributions of effective stimuli showed significant tuning differences between each interneuron type and MCs (EGC vs JGC, p = 0.37; EGC vs MC, p = 1.5e-4; JGC vs MC, p = 4.9e-5; Kolmogorov–Smirnov test; Fig. 5B, left). When naturalistic stimuli were excluded (Fig. 5B right) these effects were even more pronounced (EGC vs JGC, p = 0.063; EGC vs MC, p = 6.1e-7; JGC vs MC, p = 3.3e-5; Kolmogorov–Smirnov test). EGCs, JGCs and MCs have similar distribution pattern across monomolecular steroid stimuli (Fig. 5C). For example, A6940, P3817, Q1570 and Q3910 activated the largest number of neurons in all three cell populations, while A7010, E0893, E1050 and E4105 activated the least number of neurons. Overall, the tuning patterns observed to monomolecular sulfated steroids in all cell types were consistent with previous studies (Meeks et al., 2010; Turaga and Holy, 2012; Hammen et al., 2014).

**Figure 5.**
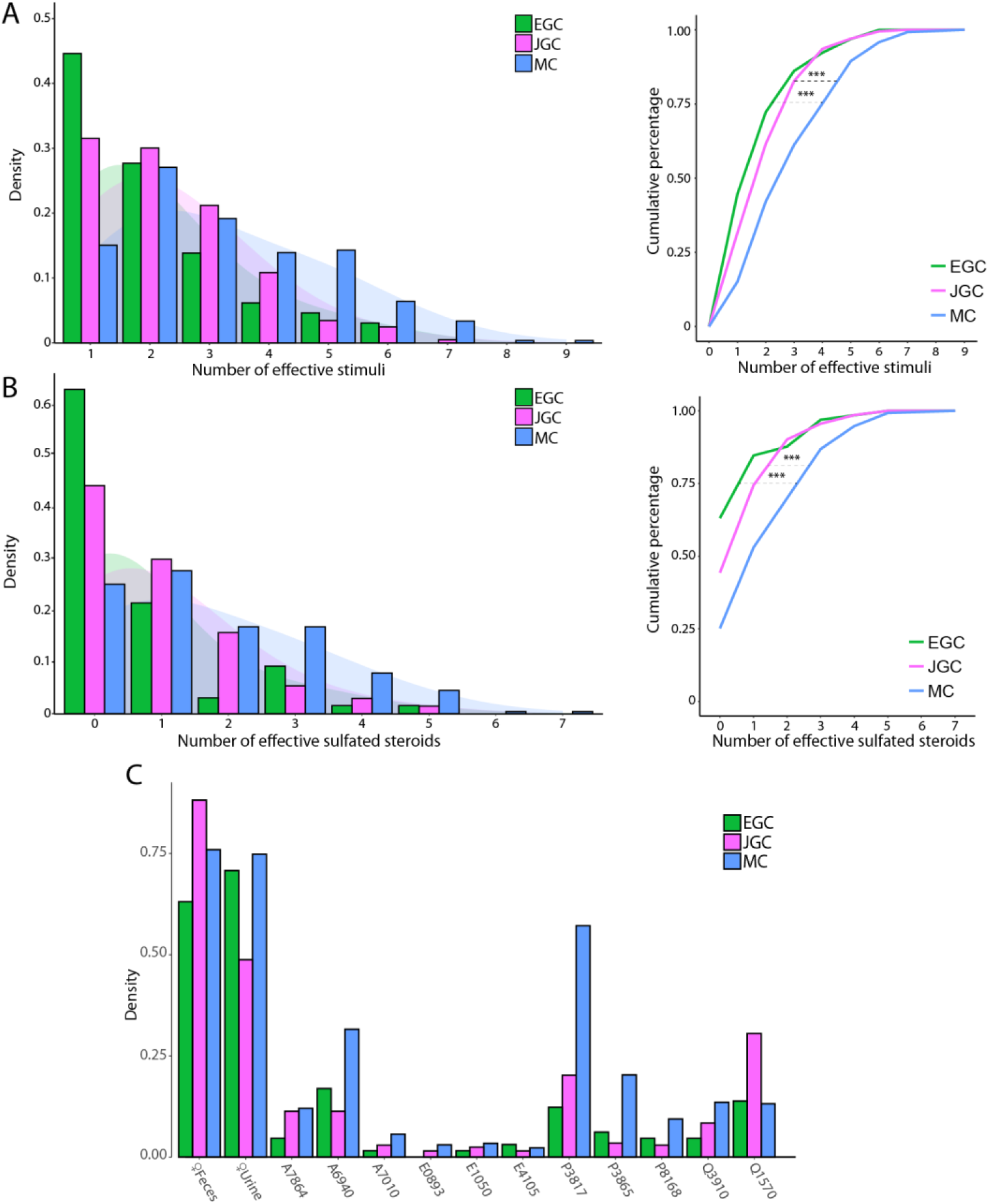
(**A**). Left: histograms of the number of effective stimuli for EGCs, JGCs, and MCs. The shaded regions indicate Gaussian kernel densities. Right: cumulative distribution of the number of effective stimuli for each cell type. EGC vs MC, p = 1.5e-4; EGC vs JGC, p = 0.37; MC vs JGC, p = 4.9e-5, K-S test. (**B**). Same as in (A), with dilute urine and feces stimuli excluded. EGC vs MC, p = 6.1e-7; JGC vs MC, p = 5.4e-5; EGC vs JGC, p = 0.063, K-S test. (**C**). Percentage of EGCs, JGCs, and MCs that responded to each stimulus in the panel.

JGC and EGC responses to natural ligand blends were overrepresented compared to MCs (Figs. 2C, 3C, 4E). Specifically, 63% of EGCs (Fig. 4E), and 44.3% of JGCs (Fig. 3C) responded exclusively to natural ligand blends, compared to 25.2% for MCs (Fig. 2C). Conversely, 9% of MCs are exclusively activated by sulfated steroids, compared to 6.2% of EGCs and 2.5% of JGCs. These differences in the proportion of responsive neurons may reflect complex network effects. However, they may more simply reflect differences in activation thresholds; previous studies indicated that MCs have higher signal/noise ratios and lower effective thresholds for activation than their VSN inputs (Meeks et al., 2010).

### EGC and MCs GCaMP6f signals differentially report electrophysiological responses

The observation that EGC chemosensory tuning is much sparser than MCs was contrary to our initial hypothesis. One possible explanation for this observation could be that EGC GCaMP6f signals more weakly reflect spiking activity than in MCs. To study spiking-GCaMP6f relationships, we performed 2-photon guided whole-cell patch clamp recordings on EGCs and MCs while recording their GCaMP6f signals in acute brain slices (Fig. 6C, D). We first found that resting membrane potentials in EGCs (– 86.0 mV ± 1.5 mV, n=13) were significantly hyperpolarized compared to MCs (−61.3± 1.0 mV, n=11), confirming earlier results (Gorin et al., 2016; Maksimova et al., 2019). In current clamp mode, standardized ramps of current injection resulted comparable maximal spike frequencies in EGCs (22-32 Hz, n = 4) and MCs (16-34 Hz, n = 5; Fig. 6A, B). We also observed that EGCs received high levels of excitatory synaptic input from MCs (Maksimova et al., 2019).

**Figure 6.**
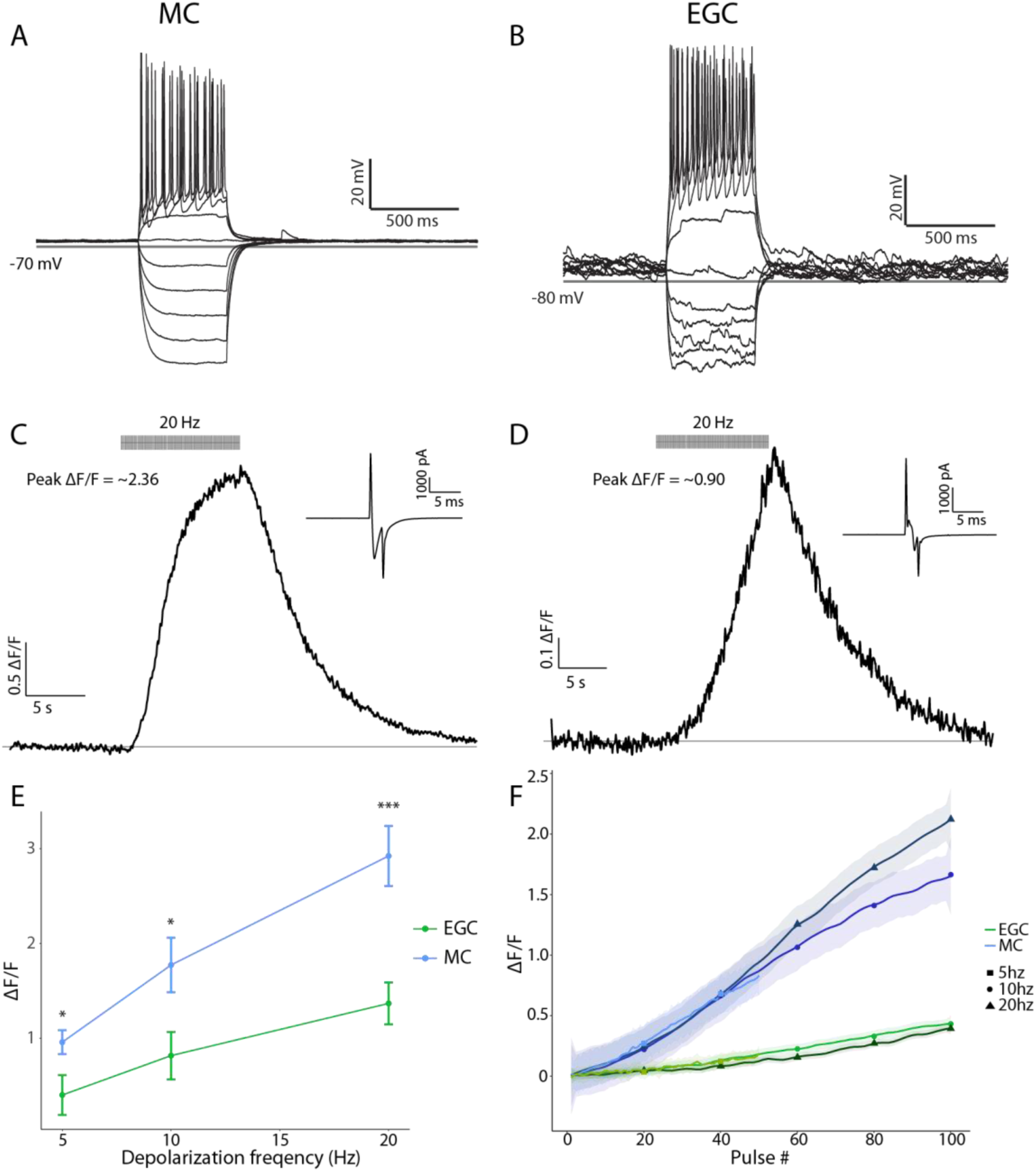
Differential GCaMP6f signaling in MCs and EGCs (**A, B**) Standardized current injection protocols elicit similar frequencies of action potentials in example MC and EGCs. (**C, D**) GCaMP6f ΔF/F trace from an example MC and EGC during a standardized 20Hz pulse train in voltage clamp mode. Insets: action currents were confirmed in response to each 2 ms step depolarization to 0 mV. (**E**) Peak ΔF/F across depolarization frequencies. 5 Hz, p = 0.045; 10 Hz, p = 0.020; 20 Hz, p = 4.4e-4 by Student’s *t*-test. (**F**) Average spiking-to-ΔF/F relationships for MCs and EGCs at 5 Hz, 10 Hz, and 20 Hz. Traces are smoothed by local polynomial regression fitting, and the shaded regions reflect 95% confidence intervals.

To probe the relationship between GCaMP6f signal and action potentials, we voltage clamped each cell at −70 mV, and then imposed a 10 s 5 Hz, 10 Hz or 20 Hz train of step depolarizations to 0 mV (MC n = 17, EGC n = 11) (Fig. 6C, D). Even prior to these depolarizations, we found that MCs showed higher baseline fluorescence compared with EGCs (see Figs. 1B, 4B). In these experiments, both MCs and EGCs responded to step depolarization with visible action currents (Fig. 6C, D). The amplitude of these currents was generally stronger in MCs, but this is expected since MC’s have a much larger soma size (MC ∼20 μm vs EGC ∼10 μm)(Larriva-Sahd, 2008) and lower input resistance (Maksimova et al., 2019). Under 20 Hz pulse train, MCs demonstrated larger peak ΔF/F (2.92 ± 0.32, n = 17) than EGCs (1.38 ± 0.22, n = 11) (Fig. 6E). Interestingly, the ΔF/F of most of MCs (15 out of 17) showed a sigmoidal stimulus-response relationship, reaching a plateau by the end of the 10 s stimulation session (Fig. 6C, also see example recording in Supplementary Video 4). In contrast, a significant fraction of EGCs (6 out of 11) did not reach a plateau in these same conditions (Fig. 6D, Supplementary Video 5). The overall relationship of spiking-to-ΔF/F across depolarization sequences showed that, across these enforced spiking frequencies, EGC GCaMP6f signals more weakly report spiking activity than GCaMP6f signals from MCs (Fig. 6F). These results suggest that a cell-type-specific difference in GCaMP6f signals contributes to the relatively sparse tuning of EGCs compared to MCs.

To evaluate the potential contribution of EGC’s weaker spiking-to-ΔF/F relationship to our initial measurements of tuning sparseness, we estimated the equivalent number of the spikes at 20 Hz associated with each neuron’s maximum ΔF/F signal. We then applied a new spiking estimate-based threshold to each cell population based on measured relationships between spiking and ΔF/F in EGCs and MCs (Fig. 7). We specifically matched the statistical criteria to those used for GCaMP6f imaging analysis, using the equivalent number of spikes needed to surpass ΔF/F cutoffs in MCs (0.1, 0.3). For MCs, the equivalent estimated spike numbers needed to achieve 0.1 and 0.3 are 10 spikes and 24 spikes, respectively (Fig. 7A, B). The estimated spike number thresholds translated to EGC ΔF/F values of 0.004 and 0.06. To avoid overestimating EGC responsivity given these low thresholds, we kept the same across-trial p-value thresholds as our initial analysis. Using this estimated spike-based threshold, EGC tuning was broader than initial measurements. This slight broadening of EGC tuning compared to MCs weakened the statistical significance between them when including both natural blends and monomolecular ligands (p = 0.090, K-S test). However, EGC sparseness to monomolecular ligands remained statistically sparser than MCs (p = 0.0069, K-S test; Fig. 7C). These results suggested that differential GCaMP6f signaling contributed to the sparse relative tuning measured in EGCs, but do not completely account for these effects.

**Figure 7.**
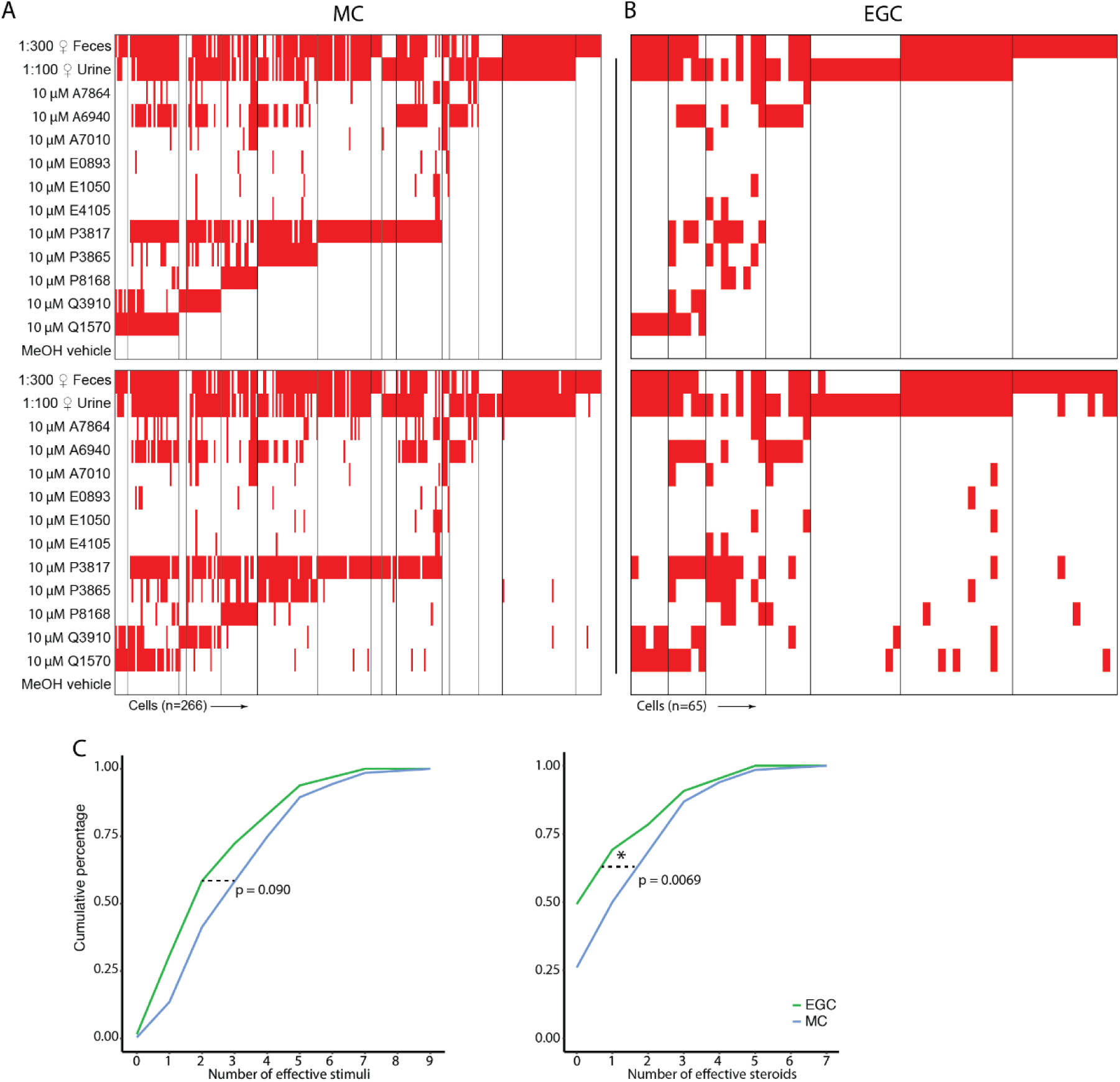
(**A, B**) Upper panels: binary responsivity heat maps for MCs (**A**) and EGCs (**B**) from GCaMP6f ΔF/F-based criteria (duplicated from Figs. 2D, 4B). Lower panels: the binary responsivity heat maps from estimated spike-based criteria. (**C**) The cumulative distributions of the total number of effective stimuli (left) and the number of effective monomolecular ligands (right) for EGCs and MCs.

### Ex vivo EGC whole cell recordings reveal broad subthreshold responsiveness

Measuring the relationships between spiking and ΔF/F in MCs and EGCs did not fully account for the sparseness of observed EGC tuning. Moreover, spiking-to-ΔF/F estimates were made using artificial spike-inducing protocols that may not reflect spiking conditions in the *ex vivo* chemosensory tuning experiments. Another possible explanation for EGC tuning sparseness could be EGCs’ highly hyperpolarized resting membrane potentials, which our data and previous studies indicated is ∼15-20 mV hyperpolarized compared to other AOB neurons (Maksimova et al., 2019). This extreme resting hyperpolarization could prevent EGC spiking in all but the strongest stimulation conditions, potentially preventing the observation of robust GCaMP6f signals. We therefore performed 2-photon fluorescence-guided whole-cell patch clamp recordings on Cort+ EGCs in the *ex vivo* preparation (Fig. 8). We performed these experiments in *Cort-2A-Cre* mice crossed to a cre-dependent tdTomato reporter line (Madisen et al., 2010), which improved our ability to identify EGC somata at rest. Using techniques similar to (Häusser and Margrie, 2013), we achieved the whole-cell configuration, then maintained each cell in current clamp near its initial resting potential via DC current injection. EGC resting membrane potentials in the *ex vivo* preparations, measured immediately after break-in, were depolarized compared to AOB slices (−63.7 ± 2.0 mV, n = 18). The reasons for the discrepancy were not clear, given that the internal and external solutions were identical to those in slice experiments. Nevertheless, we decided to maintain patched EGCs at these relatively depolarized potentials because they more likely reflected the state of EGCs in this *ex vivo* preparation (i.e. during GCaMP6f imaging experiments; Figs. 2, 4). Importantly, the observation that EGCs had relatively depolarized resting membrane potentials in the *ex vivo* preparation suggests that, if anything, EGCs in this preparation might be much closer to action potential threshold than was suggested by resting potentials measured in slice experiments.

**Figure 8.**
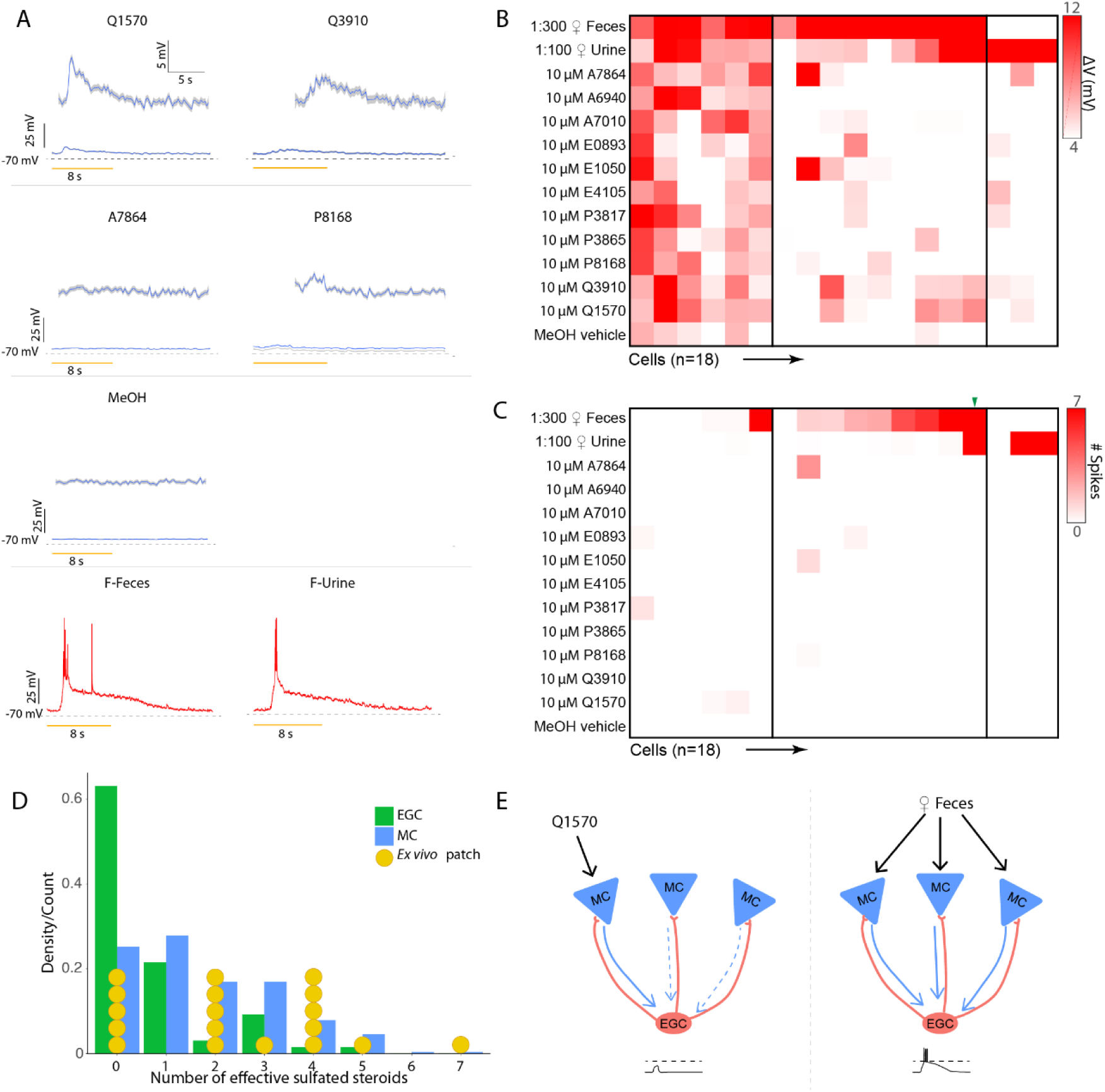
(**A**). Average voltage traces of an example EGC in the *ex vivo* preparation during VNO stimulation. Q1570, Q3910 and P8168 reliably elicit subthreshold EPSPs. Urine and fecal extracts cause action potentials. (**B**). Heatmap of average voltage changes of 18 recorded EGCs. (**C**). Heatmap of the spiking responses of the same 18 EGCs. (**D**). Comparison between responsiveness measured via Ca^2+^ imaging and patch clamp. (**E**). Threshold integration model of MC-EGC connectivity.

We recorded EGC responses while we stimulated the VNO with the same panel of chemical cues used in our GCaMP6f recordings (Fig. 8). Consistent with our GCaMP6f results, action potentials were reliably triggered in EGCs following VNO stimulation with dilute mouse urine or feces (natural ligand blends) but not by any monomolecular ligands in the stimulation panel (Fig. 8A). The spiking responses to peripheral stimulation typically included burst firing early in the stimulus presentation (11.90 ± 2.56 Hz, n = 20) followed by a long-lasting subthreshold decay period that extended well beyond the stimulation window (mean 90-10% decay time = 9.689 ± 0.436 s, n = 76 cell-stimulation pairs). The depolarization decay kinetics are comparable for naturalistic stimuli (mean 90-10% decay time = 10.255 ± 0.493 s, n = 31 cell-stimulation pairs) and monomolecular ligands (mean 90-10% decay time = 9.299 ± 0.651 s, n = 45 cell-stimulation pairs). Importantly, we readily observed broad subthreshold responses to many monomolecular steroid ligands (Fig. 8A). These subthreshold responses were tightly coupled to the onset of chemostimulation and were reliable across repeated trials (Fig. 8A). The overall envelope of depolarization was consistent with the time course of activation of MC GCaMP6f activity (Figs. 1-2) (mean 90-10% decay time = 8.085 ± 0.0644s, n = 978 cell-stimulation pairs) and previous studies (Wagner et al., 2006; Hendrickson et al., 2008; Meeks et al., 2010; Yoles-Frenkel et al., 2018).

Investigating the patterns of EGC subthreshold responsiveness revealed much broader MC integration than was indicated by GCaMP6f imaging experiments. Out of 18 responsive EGCs, 13 showed reliable subthreshold or action potential activity evoked by monomolecular sulfated steroids (Fig. 8B, p < 0.05 compared to the vehicle control, Wilcoxon rank sum test). Comparing the summed sub- and suprathreshold tuning to suprathreshold-only tuning in this group of patched cells revealed the major source of discrepancy between MC and EGC tuning (Fig. 8B, 8C). For example, 4 of the 18 recorded cells, despite clear subthreshold responses to these stimuli, did not spike at all, and presumably would have been deemed completely unresponsive in GCaMP6f imaging experiments. Of the 14 cells that spiked in response to these stimuli, the majority (9/14) spiked only in response to naturalistic ligand blends, in agreement with GCaMP6f-based results (Fig. 4D). When we included subthreshold activation into our criteria for stimulus responsivity, the distribution of EGC tuning was significantly right skewed (broader) than EGC distribution determined by the GCaMP6f imaging signal (p = 2.2e-4, Kolmogorov–Smirnov test), and became statistically indistinguishable from MCs (p = 0.21, Kolmogorov–Smirnov test; Fig. 8D). These results help to explain our *ex vivo* Ca^2+^ imaging observations and support the hypothesis that EGCs broadly integrate from MCs (Fig. 8E). However, these data also show that *Cort*+ EGCs are only effectively driven to spike by stimuli that collectively elicit spikes in a large fraction of AOB mitral cells (in this stimulus panel, dilute mouse urine and feces). Collectively, these results suggest that AOB EGCs perform fundamentally different roles in the AOB than PV-EPL interneurons play in the MOB. Though these cells are broad integrators, they are sparsely tuned at the level of spiking, suggesting that their activity is only stimulated in conditions in which a large ensemble of MCs is simultaneously activated.

## Discussion

### Cell type-specific functional studies in the AOB ex vivo preparation

Our knowledge of interneuron function in the AOB is generally lacking due to persistent technical challenges to recording from these neurons during VNO stimulation. For example, the AOB’s precarious position beneath the rhinal sinus and opposed to the prefrontal cortex creates a physical barrier for direct access to AOB neurons. The *ex vivo* preparation of the early mouse AOS, which allows optical access to the AOB while preserving VNO-AOB functional connectivity, overcomes some of the major hurdles to performing cell type-specific investigations of chemosensory tuning. The results of *ex vivo* studies do come with some limitations. For example, the *ex vivo* preparation eliminates the influence of potential feedback neuromodulation from downstream brain areas, which is clearly important for AOB circuit function (Oboti et al., 2018). Despite this significant limitation, by removing some *in vivo* complexities, the *ex vivo* approach has clear advantages for dissecting the basic structure and function of the AOB circuit.

By combining *ex vivo* methods with 2-photon microscopy and genetic tools for cell-type specific manipulation in the nervous system, we were able to perform the first studies of chemosensory tuning in genetically-defined AOB interneuron subsets. The specific cell types explored in this study, namely MCs, JGCs, and EGCs, represent 3 of the 4 major neuronal classes (with the remaining major class being the internal granule cells, or IGCs). These studies allowed us to produce quantitative comparisons of each cell type’s stimulus-response characteristics, and to do so across multiple randomized stimulus trials to reduce the possible impact of spontaneous activity (Holy et al., 2000; Nodari et al., 2008; Meeks et al., 2010). The utility of this combination of techniques for studying AOB circuit function is thus clear, and the results of these experiments allowed us to reveal key differences in the function of AOB EGCs compared to superficially-similar PV-EPL interneurons in the MOB (Kato et al., 2013; Miyamichi et al., 2013).

### AOB EGCs are broadly innervated, but sparsely tuned to chemosensory stimuli

AOB MCs are capable of integrating excitatory input from VSNs that express different sensory vomeronasal receptors (Wagner et al., 2006) and are differentially tuned to sensory input (Meeks et al., 2010). As such, observing a high amount of tuning diversity in MCs to both naturalistic stimulation and a well-characterized panel of monomolecular sulfated steroids (Fig. 2) was expected. The specific patterns of integration by MCs adds to a growing list of studies indicating that these cells support the encoding of the identity of a chemosignal-emitting animal (Luo et al., 2003; Hendrickson et al., 2008; Ben-Shaul et al., 2010; Tolokh et al., 2013). The high degree of activation of MCs by the monomolecular sulfated steroid ligands in this panel also further supports the notion that these cells possess higher coding robustness than their VSN inputs, a feature shared by principal neurons in other sensory circuits and species (Bhandawat et al., 2007; Meeks et al., 2010; Zhu et al., 2013).

Our *apriori* hypothesis was that AOB EGCs were functionally analogous to MOB PV-EPL interneurons, so we were surprised when we observed significantly sparser chemosensory tuning in EGCs compared to their upstream MCs inputs. While many EGCs were reliably activated by female BALB/c mouse urine or fecal extracts, few showed responsiveness to monomolecular ligands (Fig. 4). This result was counterintuitive, given that EGCs have extensive spinous dendritic arborizations in the ECL and receive a constant barrage of strong glutamatergic excitation from MCs, even in the absence of VSN activation (Maksimova et al., 2019). We investigated whether differential GCaMP6f signaling in EGCs and MCs could contribute to this observation. Specifically, EGC GCaMP6f signals more weakly report spiking activity than GCaMP6f signals in MCs (Fig. 6). The physiological mechanisms underlying differential GCaMP6f signaling can be difficult to experimentally pinpoint, but may include variable expression of cytosolic Ca^2+^ buffers (Schwaller, 2010), presence or absence of somatic Ca^2+^- permeant ion channels or differential activation of calcium-induced Ca^2+^ release from intracellular stores (Verkhratsky and Shmigol, 1996). Regardless of the mechanisms underlying this phenomenon, these results provide important insights into the use of GCaMP6f as an activity reporter in AOB EGCs.

Despite the differences in GCaMP6f signaling between EGCs and MCs, this effect was unable to fully explain the observed sparse tuning in EGCs (Fig. 7). EGCs were previously noted for having extremely hyperpolarized resting membrane potentials (Maksimova et al., 2019), which suggested that EGCs may possess very high thresholds for action potential generation from resting states. Whole-cell patch clamp studies of EGCs in the *ex vivo* preparation revealed that, despite using the same internal and external solutions as in slice experiments, EGC resting membrane potentials were more depolarized in the *ex vivo* preparation than in slices. This could be the result of incomplete perfusion of the relatively low [K^+^]_o_ in the aCSF (2.5 mM), or perhaps due to an overall higher excitatory tone in this preparation (or both). Importantly, these resting membrane potential measurements were made in the same conditions in which EGC GCaMP6f chemosensory tuning measurements were made, suggesting that EGC resting hyperpolarization has a less dramatic impact on tuning sparseness than expected based on slice results. In these *ex vivo* whole-cell recordings, we measured chemosensory tuning responses and observed rich subthreshold sulfated steroid-evoked activity, but little spiking except in response to stimulation with urine or feces (Fig. 8). Thus, despite mildly depolarized resting membrane potentials in the *ex vivo* preparation, EGCs demonstrate resistance to action potential generation unless a very large MC ensemble is simultaneously active (as is the case when the VNO is stimulated with mouse urine and feces). Physiological mechanisms that could contribute to high EGC thresholds for spiking could include selective expression of leak channels on EGC dendrites and shunting inhibition by other interneurons (Chamberland and Topolnik, 2012). Future studies will be needed to investigate the source of high EGC spiking thresholds.

It is worth also noting that although the *Cort-T2A-cre* transgenic mice used in these studies selectively labels AOB EGCs, the Cort+ EGC population likely represents a fraction of the total EGC population (Maksimova et al., 2019). As such, it is possible that the chemosensory tuning we observed represents a specific subgroup of Cort+ EGCs. This seems unlikely, given that no differences were found between EGC morphologies, intrinsic, and synaptic features across several transgenic lines that label these cells (Maksimova et al., 2019). Also, even though *Cort+* EGC labeling spans the anterior AOB (which is sensitive to sulfated steroids and many urinary and fecal cues) and posterior AOB (sensitive to urinary proteins and exocrine gland-secreted peptides), our optical recordings were largely confined to portions of the anterior AOB where responsiveness to the cues in our panel is most prominent (Meeks et al., 2010; Doyle et al., 2014). As such, it may be the case that EGCs in the posterior AOB have different tuning qualities than is indicated in this study.

### Implications for models of AOB information processing

The complex physiological properties of EGCs and their chemosensory tuning are becoming clearer, but the impacts of EGC activation on MC function remain unclear. Many EGCs are labeled in *Gad2-IRES-cre* transgenic lines (Maksimova et al., 2019), consistent with a GABAergic phenotype. EGCs are axonless and have a spinous dendrites that closely appose MC dendrites, and AOB MCs are known to form reciprocal dendro-dendritic synapses with other AOB interneurons (Jia et al., 1999; Taniguchi and Kaba, 2001; Castro et al., 2007; Larriva-Sahd, 2008). Seemingly analogous PV-EPL interneurons in the MOB have been shown to be broadly inhibitory (Kato et al., 2013; Miyamichi et al., 2013). All of these pieces of evidence point to a reciprocal inhibitory function for EGCs, and future studies will be able to further elucidate EGCs’ impact on MC function and information flow through the AOB.

Regardless of the specific mechanisms underlying EGCs’ specific chemosensory tuning features, the observation that EGCs rarely spike in the absence of a naturalistic ligand blend has important implications for AOB circuit function. First, these results suggest that EGCs have unique roles in AOB processing that are different from PV-EPL interneurons in MOB. Despite their superficially similar circuit architectures, seemingly analogous neural types in the MOB and AOB have repeatedly been shown to have fundamentally different physiological properties (Shipley and Adamek, 1984; Jia et al., 1999; Araneda and Firestein, 2006; Wagner et al., 2006; Castro et al., 2007; Smith et al., 2015). In the MOB, PV-EPL interneurons are activated by many monomolecular odorants with low thresholds, resulting in chemosensory tuning that is close to a simple linear addition of input MCs’ tuning maps (Kato et al., 2013). This quality benefits unbiased monitoring of MC activity and supports divisive normalization of MCs based on the overall population response (Kato et al., 2013; Miyamichi et al., 2013). In contrast, AOB EGCs appear to have extremely high effective thresholds despite receiving synaptic input from many MCs, which may strongly bias their activity away from monomolecular ligands and towards the blends of ligands found in natural excretions (Nodari et al., 2008). Since natural vomeronasal social cues are only known to exist in the form of such complex blends, the difference in monomolecular tuning sparseness between AOB EGCs and MOB PV-EPL interneurons and AOB EGCs might reflect macroscopic differences in the natural statistics of ligand sampling between these two chemosensory pathways. It may also be the case that MC inhibition by EGCs takes place locally at reciprocal dendrodendritic synapses in a spiking-independent manner, as has been observed in the MOB (Isaacson and Strowbridge, 1998; Schoppa et al., 1998; Chen et al., 2000; Halabisky et al., 2000; Isaacson, 2001; Egger et al., 2005; Bywalez et al., 2015; Lage-Rupprecht et al., 2018).

AOB EGCs thus appear to be, at a minimum, operating in a manner that is strikingly different than PV-EPL interneurons in the MOB, and tend not to be strongly active in the absence of broad AOB activation, raising questions about their *in vivo* roles in sensory processing. Several studies have reported individual vomeronasal ligands capable of evoking significant behavioral effects through the AOS (Chamero et al., 2007; Haga et al., 2010; Papes et al., 2010). If our results hold for the ligands that drive these behaviors, it seems unlikely that these particular chemosensory exposures engage EGCs, which may have important implications for information flow through the AOB towards its downstream targets in the limbic system (Martinez-Marcos, 2009; Gutiérrez-Castellanos et al., 2014; Stowers and Liberles, 2016). In sum, these experiments contribute a wealth of information about chemosensory tuning of specific AOB cell types, adding important quantitative constraints on the role of inhibitory interneurons on AOB circuit function.

## Acknowledgements

We thank Jie Cao, Marina Maksimova, Daniel Dinh, Natasha Browder, and Cara Nielson for methodological advice and technical support.

## References

Araneda RC, Firestein S (2006) Adrenergic Enhancement of Inhibitory Transmission in the Accessory Olfactory Bulb. The Journal of Neuroscience 26:3292–3298.

Ben-Shaul Y, Katz LC, Mooney R, Dulac C (2010) In vivo vomeronasal stimulation reveals sensory encoding of conspecific and allospecific cues by the mouse accessory olfactory bulb. Proceedings of the National Academy of Sciences 107:5172–5177.

Bhandawat V, Olsen SR, Gouwens NW, Schlief ML, Wilson RI (2007) Sensory processing in the Drosophila antennal lobe increases reliability and separability of ensemble odor representations. Nature Neuroscience 10:1474–1482.

Braganza O, Beck H (2018) The Circuit Motif as a Conceptual Tool for Multilevel Neuroscience. Trends in Neurosciences 41:128–136.

Brennan PA (2001) The vomeronasal system. Cellular and Molecular Life Sciences CMLS 58:546–555.

Brennan PA, Keverne EB (1997) Neural mechanisms of mammalian olfactory learning. Progress in Neurobiology 51:457–481.

Brennan PA, Binns EK (2005) Vomeronasal mechanisms of mate recognition in mice. Chemical senses.

Brennan PA, Kendrick KM, Neuroscience K-EB (1995) Neurotransmitter release in the accessory olfactory bulb during and after the formation of an olfactory memory in mice. Neuroscience.

Bywalez WG, Patirniche D, Rupprecht V, Stemmler M, Herz A, Pálfi D, Rózsa B, Egger V (2015) Local Postsynaptic Voltage-Gated Sodium Channel Activation in Dendritic Spines of Olfactory Bulb Granule Cells. Neuron 85:590–601.

Cansler HL, Maksimova MA, Meeks JP (2017) Experience-dependent plasticity in accessory olfactory bulb interneurons following male-male social interaction. bioRxiv:127589.

Castro JB, Hovis KR, Urban NN (2007) Recurrent Dendrodendritic Inhibition of Accessory Olfactory Bulb Mitral Cells Requires Activation of Group I Metabotropic Glutamate Receptors. The Journal of Neuroscience 27:5664–5671.

Chamberland S, Topolnik L (2012) Inhibitory control of hippocampal inhibitory neurons. Frontiers in Neuroscience 6:165.

Chamero P, Marton TF, Logan DW, Flanagan K, Cruz JR, Saghatelian A, Cravatt BF, Stowers L (2007) Identification of protein pheromones that promote aggressive behaviour. Nature 450:899.

Chen TW, Wardill TJ, Sun Y, Pulver SR, Renninger SL, Baohan A, Schreiter ER, Kerr RA, Orger MB, Jayaraman V, Looger LL, Svoboda K, Kim DS (2013) Ultrasensitive fluorescent proteins for imaging neuronal activity. Nature 499:295–300.

Chen WR, Xiong W, Shepherd GM (2000) Analysis of Relations between NMDA Receptors and GABA Release at Olfactory Bulb Reciprocal Synapses. Neuron 25:625–633.

Doyle WI, Meeks JP (2017) Heterogeneous effects of noradrenaline on spontaneous and stimulus-driven activity in the male accessory olfactory bulb. Journal of neurophysiology 117.

Doyle WI, Hammen GF, Meeks JP (2014) *Ex Vivo* Preparations of the Intact Vomeronasal Organ and Accessory Olfactory Bulb. Journal of Visualized Experiments.

Doyle WI, Dinser JA, Cansler HL, Zhang X, Dinh DD, Browder NS, Riddington IM, Meeks JP (2016) Faecal bile acids are natural ligands of the mouse accessory olfactory system. Nature Communications 7:11936.

Egger V, Svoboda K, Mainen ZF (2005) Dendrodendritic Synaptic Signals in Olfactory Bulb Granule Cells: Local Spine Boost and Global Low-Threshold Spike. The Journal of Neuroscience 25:3521–3530.

Gao Y, Budlong C, Durlacher E, Davison IG (2017) Neural mechanisms of social learning in the female mouse. eLife 6.

Geramita M, Urban NN (2017) Differences in Glomerular-Layer-Mediated Feedforward Inhibition onto Mitral and Tufted Cells Lead to Distinct Modes of Intensity Coding. Journal of Neuroscience 37:1428–1438.

Gorin M, Tsitoura C, Kahan A, Watznauer K, Drose DR, Arts M, Mathar R, O’Connor S, Hanganu-Opatz IL, Ben-Shaul Y, Spehr M (2016) Interdependent Conductances Drive Infraslow Intrinsic Rhythmogenesis in a Subset of Accessory Olfactory Bulb Projection Neurons. The Journal of Neuroscience 36:3127–3144.

Gutiérrez-Castellanos N, Pardo-Bellver C, Martínez-García F, Lanuza E (2014) The vomeronasal cortex – afferent and efferent projections of the posteromedial cortical nucleus of the amygdala in mice. European Journal of Neuroscience 39:141–158.

Haga S, Hattori T, Sato T, Sato K, Matsuda S, Kobayakawa R, Sakano H, Yoshihara Y, Kikusui T, Touhara K (2010) The male mouse pheromone ESP1 enhances female sexual receptive behaviour through a specific vomeronasal receptor. Nature 466:118.

Halabisky B, Friedman D, Radojicic M, Strowbridge BW (2000) Calcium Influx through NMDA Receptors Directly Evokes GABA Release in Olfactory Bulb Granule Cells. Journal of Neuroscience 20:5124–5134.

Hammen GF, Turaga D, Holy TE, Meeks JP (2014) Functional organization of glomerular maps in the mouse accessory olfactory bulb. Nat Neurosci 17.

Häusser M, Margrie TW (2013) Two-Photon Targeted Patching and Electroporation In Vivo. Cold Spring Harbor Protocols 2014.

Hendrickson RC, Krauthamer S, Essenberg JM, Holy TE (2008) Inhibition shapes sex selectivity in the mouse accessory olfactory bulb. J Neurosci 28.

Holy TE, Dulac C, Meister M (2000) Responses of Vomeronasal Neurons to Natural Stimuli. Science 289:1569–1572.

Isaacson JS (2001) Mechanisms governing dendritic γ-aminobutyric acid (GABA) release in the rat olfactory bulb. Proceedings of the National Academy of Sciences 98:337–342.

Isaacson JS, Strowbridge BW (1998) Olfactory Reciprocal Synapses: Dendritic Signaling in the CNS. Neuron 20:749–761.

Jeanne JM, Wilson RI (2015) Convergence, Divergence, and Reconvergence in a Feedforward Network Improves Neural Speed and Accuracy. Neuron 88.

Jia C, Chen WR, Shepherd GM (1999) Synaptic Organization and Neurotransmitters in the Rat Accessory Olfactory Bulb. Journal of Neurophysiology 81:345–355.

Kaba H, Keverne EB (1988) The effect of microinfusions of drugs into the accessory olfactory bulb on the olfactory block to pregnancy. Neuroscience 25:1007–1011.

Kaba H, Nakanishi S (1995) Synaptic Mechanisms of Olfactory Recognition Memory. Rev Neuroscience 6.

Kaba H, Huang G-ZZ (2005) Long-term potentiation in the accessory olfactory bulb: a mechanism for olfactory learning. Chem Senses 30 Suppl 1.

Kato HK, Gillet SN, Peters AJ, Isaacson JS, Komiyama T (2013) Parvalbumin-expressing interneurons linearly control olfactory bulb output. Neuron 80.

Lage-Rupprecht V, Jodar T, Yeghiazaryan G, Rozsa B, Egger V (2018) Local reciprocal release of GABA from dendritic spines of olfactory bulb granule cells requires local sodium channel activation and occurs on both fast and slow timescales. bioRxiv:440198.

Larriva-Sahd J (2008) The accessory olfactory bulb in the adult rat: a cytological study of its cell types, neuropil, neuronal modules, and interactions with the main olfactory system. J Comp Neurol 510:309–350.

Luo M, Fee MS, Katz LC (2003) Encoding Pheromonal Signals in the Accessory Olfactory Bulb of Behaving Mice. Science 299:1196–1201.

Madisen L, Zwingman TA, Sunkin SM, Oh SW, Zariwala HA, Gu H, Ng LL, Palmiter RD, Hawrylycz MJ, Jones AR, Lein ES, Zeng H (2010) A robust and high-throughput Cre reporting and characterization system for the whole mouse brain. Nature neuroscience 13:133–140.

Maksimova MA, Cansler HL, Zuk KE, Torres JM, Roberts DJ, Meeks JP (2019) Interneuron functional diversity in the mouse accessory olfactory bulb. bioRxiv:552463.

Martinez-Marcos A (2009) On the organization of olfactory and vomeronasal cortices. Progress in Neurobiology 87:21–30.

Matsuoka M, Kaba H, of … M-K (2004) Remodeling of reciprocal synapses associated with persistence of long-term memory. European Journal of ….

Matsuoka M, Kaba H, Mori Y, Neuroreport I-M (1997) Synaptic plasticity in olfactory memory formation in female mice. Neuroreport.

Meeks JP, Holy TE (2009) An *ex vivo* preparation of the intact mouse vomeronasal organ and accessory olfactory bulb. Journal of Neuroscience Methods 177:440–447.

Meeks JP, Arnson HA, Holy TE (2010) Representation and transformation of sensory information in the mouse accessory olfactory system. Nature neuroscience.

Miyamichi K, Shlomai-Fuchs Y, Shu M, Weissbourd BC, Luo L, Mizrahi A (2013) Dissecting local circuits: parvalbumin interneurons underlie broad feedback control of olfactory bulb output. Neuron 80.

Mohrhardt J, Nagel M, Fleck D, Ben-Shaul Y, Spehr M (2018) Signal Detection and Coding in the Accessory Olfactory System. Chemical Senses 43:667–695.

Moriya-Ito K, Endoh K, Fujiwara-Tsukamoto Y, Ichikawa M (2013) Three-dimensional reconstruction of electron micrographs reveals intrabulbar circuit differences between accessory and main olfactory bulbs. Frontiers in Neuroanatomy 7:5.

Nagai Y, Sano H, Yokoi M (2005) Transgenic expression of Cre recombinase in mitral/tufted cells of the olfactory bulb. genesis 43:12–16.

Nodari F, Hsu F-F, Fu X, Holekamp TF, Kao L-F, Turk J, Holy TE (2008) Sulfated Steroids as Natural Ligands of Mouse Pheromone-Sensing Neurons. The Journal of Neuroscience 28:6407–6418.

Oboti L, Russo E, Tran T, Durstewitz D, Corbin JG (2018) Amygdala Corticofugal Input Shapes Mitral Cell Responses in the Accessory Olfactory Bulb. eNeuro 5:18.

Papes F, Logan DW, Stowers L (2010) The Vomeronasal Organ Mediates Interspecies Defensive Behaviors through Detection of Protein Pheromone Homologs. Cell 141:692–703.

Rothermel M, Brunert D, Zabawa C, Díaz-Quesada M, Wachowiak M (2013) Transgene Expression in Target-Defined Neuron Populations Mediated by Retrograde Infection with Adeno-Associated Viral Vectors. The Journal of Neuroscience 33:15195–15206.

Schoppa NE, Kinzie MJ, Sahara Y, Segerson TP, Westbrook GL (1998) Dendrodendritic Inhibition in the Olfactory Bulb Is Driven by NMDA Receptors. Journal of Neuroscience 18:6790–6802.

Schwaller B (2010) Cytosolic Ca2+ Buffers. Cold Spring Harbor Perspectives in Biology 2.

Shipley MT, Adamek GD (1984) the connections of the mouse olfactory bulb: A study using orthograde and retrograde transport of wheat germ agglutinin conjugated to horseradish peroxidase. Brain Research Bulletin 12:669–688.

Smith RS, Hu R, DeSouza A, Eberly CL, Krahe K, Chan W, Araneda RC (2015) Differential Muscarinic Modulation in the Olfactory Bulb. The Journal of Neuroscience 35:10773–10785.

Stowers L, Liberles SD (2016) State-dependent responses to sex pheromones in mouse. Current Opinion in Neurobiology 38:74–79.

Su C-Y, Menuz K, Carlson JR (2009) Olfactory perception: receptors, cells, and circuits. Cell 139:45–59.

Taniguchi H, He M, Wu P, Kim S, Paik R, Sugino K, Kvitsiani D, Kvitsani D, Fu Y, Lu J, Lin Y, Miyoshi G, Shima Y, Fishell G, Nelson SB, Huang ZJ (2011) A resource of Cre driver lines for genetic targeting of GABAergic neurons in cerebral cortex. Neuron 71.

Taniguchi M, Kaba H (2001) Properties of reciprocal synapses in the mouse accessory olfactory bulb. Neuroscience 108:365–370.

Tolokh, II, Fu X, Holy TE (2013) Reliable sex and strain discrimination in the mouse vomeronasal organ and accessory olfactory bulb. J Neurosci 33:13903–13913.

Turaga D, Holy TE (2012) Organization of vomeronasal sensory coding revealed by fast volumetric calcium imaging. J Neurosci 32:1612–1621.

Verkhratsky A, Shmigol A (1996) Calcium-induced calcium release in neurones. Cell Calcium 19:1–14.

Wagner S, Gresser AL, Torello AT, Dulac C (2006) A multireceptor genetic approach uncovers an ordered integration of VNO sensory inputs in the accessory olfactory bulb. Neuron 50.

Wilson NR, Runyan CA, Wang FL, Sur M (2012) Division and subtraction by distinct cortical inhibitory networks in vivo. Nature 488.

Yoles-Frenkel M, Kahan A, Ben-Shaul Y (2018) Temporal Response Properties of Accessory Olfactory Bulb Neurons: Limitations and Opportunities for Decoding. The Journal of neuroscience: the official journal of the Society for Neuroscience 38:4957–4976.

Zhu P, Frank T, Friedrich RW (2013) Equalization of odor representations by a network of electrically coupled inhibitory interneurons. Nature Neuroscience 16:1678–1686.

